# Smc3 acetylation, Pds5 and Scc2 control the translocase activity that establishes cohesin dependent chromatin loops

**DOI:** 10.1101/2021.04.21.440823

**Authors:** Nathalie Bastié, Christophe Chapard, Lise Dauban, Olivier Gadal, Frederic Beckouёt, Romain Koszul

## Abstract

Chromosome spatial organization and dynamics influence DNA-related metabolic processes. SMC complexes like cohesin are essential instruments of chromosome folding. Cohesin-dependent chromatin loops bring together distal loci to regulate gene transcription, DNA repair and V(D)J recombination processes. Here we characterize further the roles of members of the cohesin holocomplex in regulating chromatin loop expansion, showing that Scc2, which stimulates cohesin ATPase activity, is essential for the translocation process required to extend DNA loop length. Eco1-dependent acetylation of Smc3 during S phase counteracts this activity through the stabilization of Pds5, to finely tune loop sizes and stability during G2. Inhibiting Pds5 in G2 leads to a strong enlargement of pre-established, stable DNA loops, in a Scc2-dependent manner. Altogether, the study strongly supports a Scc2-mediated translocation process driving expansion of DNA loops in living cells.

## INTRODUCTION

The three-dimensional folding of chromosomes influences or modulates a number of DNA-related metabolic processes, such as gene expression, replication, repair or segregation (Marchal et al., 2019; Zheng and Xie, 2019). The development of high-throughput chromosome conformation capture techniques (e.g. Hi-C) and high-resolution imaging approaches that quantify contact frequencies and physical distances between distant DNA loci have now unveiled the multi-layered organization of genomes (Dekker and Mirny, 2016; Szabo et al., 2019). A variety of mega-base (Mb) and sub-Mb structures, such as chromatin loops or self-interacting domains, have been described along the chromosomes of all studied organisms so far including bacteria, archaea, yeast and mammals (Cockram et al., 2021; Dauban et al., 2020; Dekker and Mirny, 2016; Gibcus et al., 2018; Le et al., 2013; Marbouty et al., 2015; Muller et al., 2018; Nora et al., 2012; Szabo et al., 2019).

Multiple studies show that cohesin, a member of the structural maintenance of chromosomes (SMC) protein family well-known for its role in maintaining sister-chromatid cohesion (SCC) during mitosis (Nasmyth and Haering, 2009), is a major player in eukaryotic chromatin organisation (Dauban et al., 2020; Haarhuis et al., 2017; Lazar-Stefanita et al., 2017; Merkenschlager and Nora, 2016; Rao et al., 2017; Schalbetter et al., 2017; Wutz et al., 2020; Wutz et al., 2017). The cohesin complex consists of two Smc proteins, Smc1 and Smc3, that dimerize via their hinge domains and interact with the kleisin Scc1 via their ATPase head domains (Nasmyth and Haering, 2009). This tripartite complex forms a large ring that can accommodate one or two DNA molecules (Haering et al., 2008). Many proteins regulate the association of the cohesin holocomplex with DNA: on the one hand, Scc2 and Scc4 are involved in loading the cohesin ring on DNA; on the other hand, Wpl1 and Pds5 play a role in its release (Beckouet et al., 2016; Chan et al., 2013; Chan et al., 2012; Fernius et al., 2013; Kikuchi et al., 2016; Makrantoni and Marston, 2018). During replication, sister chromatids (SCs) are thought to be topologically entrapped within cohesin rings. To stabilize SCC from S to M phase, the releasing activity (Wpl1/Pds5) is repressed during S phase by Eco1 mediated acetylation of a pair of conserved lysine residues (K112/113) within the Smc3 ATPase head (Chan et al., 2012; Rolef Ben-Shahar et al., 2008; Rowland et al., 2009; Unal et al., 2008; Zhang et al., 2008). Smc3 acetylation is maintained throughout the G2 and M stages and only removed at the onset of anaphase by the deacetylase Hos1, following the cleavage of Scc1 by Separase (Beckouet et al., 2010; Nasmyth and Haering, 2009). Among all the proteins that regulate cohesin functions Pds5 is one of the most peculiar, in addition of being implicated in releasing activity, Pds5 promotes Smc3 acetylation by Eco1 and subsequently prevents Smc3 de-acetylation from Hos1 (Chan et al., 2013)

Recently, cohesins were shown to promote large chromatin loops along mammalian chromosomes, modulating contacts between pairs of loci sometimes separated by hundreds of kb. These loops contribute to the segmentation of interphase chromosomes in self-interacting topologically associating domains or TADs (reviewed in (Dekker and Mirny, 2016; Rowley and Corces, 2018; Yu and Ren, 2017). TADs boundaries coincide with the binding sites of transcriptional repressor CTCF in convergent orientation (Rao et al., 2014), frequently enriched in cohesin deposition. These observations suggested that cohesin organises chromatin by capturing small loops of DNA and gradually enlarging them into large structures, through an active process dubbed loop extrusion (Goloborodko et al., 2016; Nasmyth, 2001; Sanborn et al., 2015), until they encounter a roadblock corresponding to an occupied CTCF site. Experimental support for loop extrusion recently came from *in vitro* single molecule experiments showing that human cohesin (and the SMC homolog condensin) share the capability to extrude DNA (Davidson et al., 2019; Ganji et al., 2018; Kim et al., 2019). Nipbl, the mammalian homolog of Scc2, and ATP hydrolysis are essential to promote both DNA loop expansion and DNA loop maintenance *in vitro*. Scc2 is furthermore essential to stimulate cohesin’s ATPase activity (Davidson et al., 2019; Petela et al., 2018). By competing with Scc2 for binding to cohesin’s kleisin (Kikuchi et al., 2016) and with the help from Eco1, Pds5 may in addition inhibits the putative Scc2 mediated translocation process.

Structural findings further show that CTCF does not simply behaves as a passive physical boundary to cohesin extruding DNA, but suppresses the release of cohesin from DNA by competing with Wpl1 for binding on cohesin. Therefore, CTCF stabilises cohesin, and consequently chromatin DNA loops, along the chromosome (Li et al., 2020). It has been proposed that loop extrusion favours contacts between promoters and enhancers, whose contacts depend on CTCF and cohesin (Merkenschlager and Nora, 2016; Yatskevich et al., 2019). While cohesin removal has no significant impact on global gene expression (Rao et al., 2017), mutations in genes encoding the cohesin subunits have a local effect on the regulation of transcription (Merkenschlager and Nora, 2016). Recent studies further provide strong evidence for the importance of loop extrusion such as in DNA repair (Arnould et al., 2021; Piazza et al., 2020) or V(D)J recombination processes (Ba et al., 2020; Dai et al., 2021; Zhang et al., 2019a; Zhang et al., 2019b).

We and others recently showed that cohesin dependent loops are also found along yeast mitotic chromosomes (Costantino et al., 2020; Dauban et al., 2020; Paldi et al., 2020). Those loops can form independently of replication and the presence of a sister chromatid (and SCC)(Dauban et al 2020). Although CTCF is not present in this organism, and therefore cannot contribute to the definition of loop basis, the regulation of cohesin loop expansion is highly conserved between species. Notably, Pds5 and Eco1 negatively regulate DNA loop enlargement in the budding yeast as well as in mammals (Dauban et al., 2020; Wutz et al., 2020; Wutz et al., 2017). Despite important progresses in the last few years, and although the mechanisms mediating cohesin-dependent SCC are well described, how the same complex organises the folding of chromatin in *cis* remains unclear. Loop organized mitotic chromosomes also raise questions regarding the interplay between the cohesin complexes involved in SCC, and the ones involved in *cis* loop formation. Questions also relate to the elements positioning and driving loop expansion.

We show that replacing Smc3 lysine 112 and 113 residues by arginine in yeast reduces the number of mitotic DNA loops and increases the length of cis contacts along chromosome arms. We further show that Smc3 acetylation by Eco1 controls DNA loop border positions and sizes during S phase. This establishment is restricted to S phase, and depends on Pds5, which is therefore necessary to block the translocation process. Pds5 stabilizes DNA loops along the chromosomes. In addition, the loss of Scc2 activity in G2/M does not destabilize a collection of established DNA loops. Instead, wild-type Scc2 activity promotes contacts between distant loci along chromosome arms in the absence of Pds5 regulation. Scc2 is therefore required for the translocation process, in addition to its canonical role in cohesin loading.

## RESULTS

### Smc3 acetylation on K112 and K113 suppresses DNA loop expansion

The mechanisms by which Pds5 and Eco1 inhibit the expansion of cohesin-dependent chromatin loops along mitotic budding yeast chromosomes remains unknown (Dauban et al., 2020). Since Smc3 acetylation by Eco1 is essential for SCC during S phase, we wondered whether the Eco1 dependent inhibition of DNA loop expansion could also depend on its Smc3 acetylation activity. We therefore characterized the formation of DNA loops in haploid strains ectopically expressing either a non-acetylable version of Smc3 (*smc3*(*K112-113R*), here referred to as *smc3-RR*) or a wild type allele of *SMC3* integrated at LEU2, along with an auxin degradable endogenous *SMC3* (*smc3-AID*). The endogenous SMC3 is used to sustain cell viability of *smc3-RR* otherwise lethal. The effect of *smc3-RR* on chromatin folding was scrutinized in cells depleted for the endogenous Smc3 before replication and arrested in metaphase by the depletion of the APC activator Cdc20 (Methods, Figure 1A). The metaphase arrest and the depletion of Smc3-AID consecutive to auxin addition were confirmed by flow cytometry and western blot respectively (Figure 1B and 1C). Three experimental conditions can then be analyzed: the first one in which no Smc3 is present in the cell, the second in which WT Smc3 is expressed and finally, cells in which only Smc3-RR is expressed. Hi-C contact maps were generated for these three conditions in parallel to wild type and Eco1 depleted cells in metaphase (Figure 1D). The normalized contact maps and the contact probabilities as a function of genomic distance curves *P(s*) and their derivatives show that *smc3-RR* is similar to Eco1 depletion, with cis contacts between pairs of loci spreading to longer distances (> 40kb) compared to wild type cells (Figure 1D and 1E). Chromatin loops along the chromosomes (Figure 1D) were called on contact maps using the program Chromosight (Matthey-Doret et al., 2020), revealing that the number of distinguishable and visible loops along chromosome arms in the presence of *smc3-RR* (88 loops) is reduced compared to a control strain expressing an ectopic copy of SMC3 (*SMC3::LEU2*, (202 loops)) or *WT* cells (223 loops, Figure 1D). This number is similar to the number of loops detected in the absence of Eco1 (60 loops, Figure 1D). Comparison of mean loop scores (MLS; i.e. the correlation scores of a set of input coordinates with a generic kernel; Methods; Matthey-Doret et al., 2020) confirmed the reduction of the amount of loops detected in cells expressing *smc3-RR* (MLS = 0,14) or depleted for Eco1 (0,16) compared to Smc3 (0,38) or *WT* loops (0,42) (Figure 1G). Interestingly, the few detected DNA loops have their basis separated by longer distances (up to 80 kb in presence of *smc3-RR*) compared to control cells (< 50kb), as shown by the loop spectrum (Figure 1F). Altogether both results obtained with *smc3-RR* and Eco1 depletion show that the Eco1 mediated acetylation of Smc3 K112 and K113 inhibits DNA loop expansion. In the absence of Eco1 mediating Smc3 acetylation, most “stop” sites appear now bypassed, with a few hotspots standing out where most loop signal accumulate. Therefore, in addition to its role in sister chromatid-cohesion maintenance, Smc3 acetylation plays a role in stabilizing cohesin-dependent cis-contacts on DNA.

**Figure 1:**
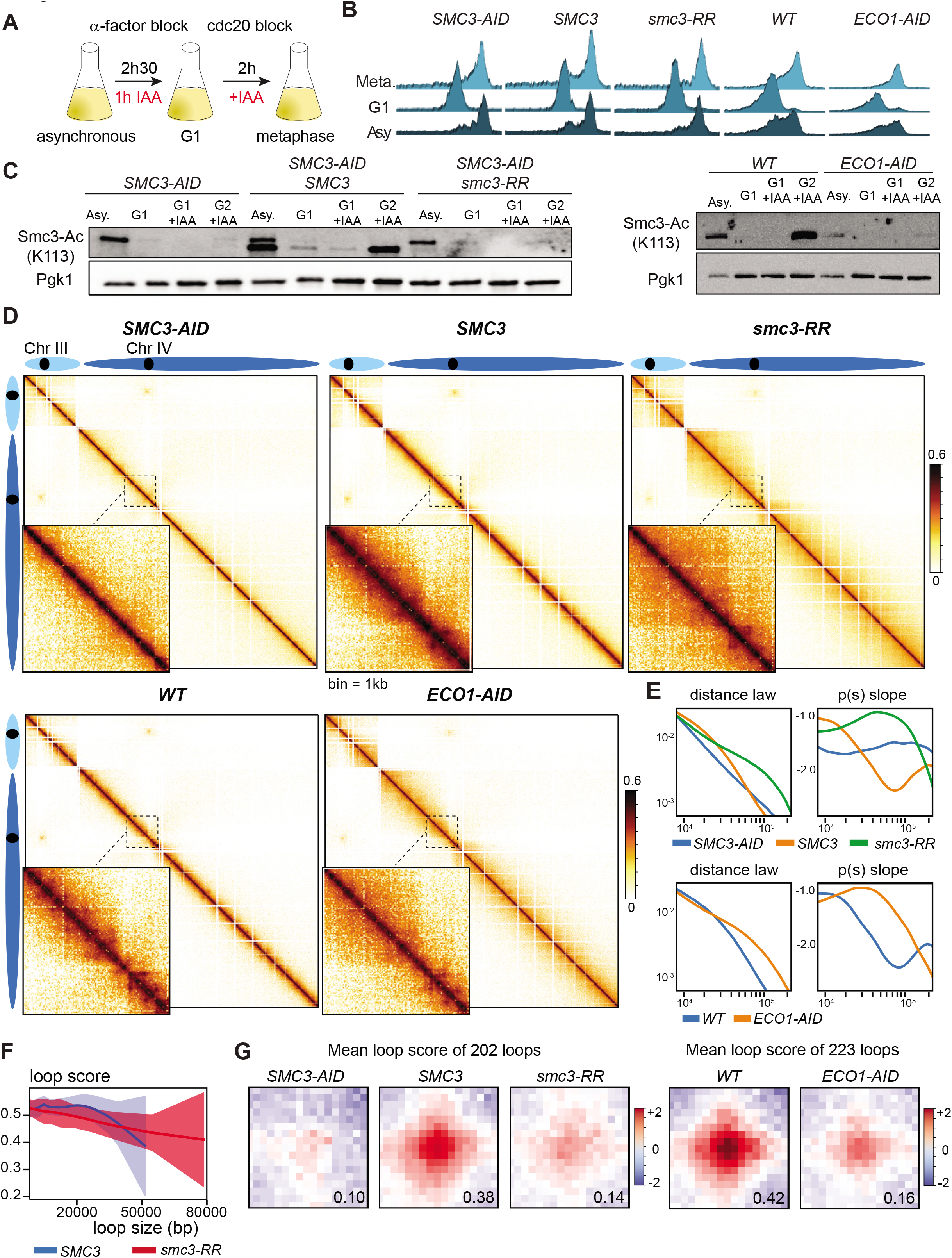
Smc3 acetylation counteracts DNA loop expansion. A. Illustration of the experimental protocol used to process yeast cells from G1 to metaphase in the presence of Smc3-RR or in the absence of Eco1-AID. B. Cell synchronisation was monitored by flow cytometry. C. Western blot assessing loss of Smc3 acetylation induced by Smc3-AID or Eco1-AID depletion. Right panel: Smc3-AID is represented by the top band while Smc3 WT is the lowest band. Pgk1 was used as loading control. D. Contact maps of chromosomes III and IV (bin 1kb) from metaphase arrested cells. Upper panel: contact map for cells without (*SMC3-AID* +IAA, left, FB134-16C) or with Smc3 (*SMC3* +IAA, middle, yNB54-4) and for cells with Smc3-RR (*smc3-RR* +IAA, right, yNB54-3). Lower panel: contact maps for cells with (WT +IAA, left, yLD116-1a) or without Eco1 (*ECO1-AID* +IAA, right, FB133-20c). Chromatin interactions along a region (300kb to 510kb) of the chromosome IV are represented in the black square. E. Contact probability curves Pc(s) representing contact probability as a function of genomic distance (bp) and their respective derivative curves F. Loop spectrum indicating scores in function of loop size. G. Mean profile heatmap of loops called by chromosight.

### Eco1 is not required during mitosis to inhibit DNA loop expansion on yeast chromosomes

To assess whether Eco1 activity in post replicative cells is required to inhibit DNA loops expansion, we degraded Eco1-AID from cells blocked in G2/M by cdc20 depletion (Methods; Figure 2A, 2B and 2C). Once arrested in metaphase, and upon auxin addition, chromatin contacts were analysed using Hi-C. Control samples without auxin addition were processed in parallel. Normalized Hi-C maps and the corresponding *P*(*s*) curves reveal that Eco1 depletion from post-replicative cells has no substantial consequences on chromatin contacts (Figure 2D and 2E). Note that the AID epitope appears to slightly modify Eco1 function during S phase, as cells expressing Eco1-AID present slightly longer mitotic loop compared to wild type Eco1 cells. Loop calling using Chromosight further showed that degradation of Eco1 during G2/M does not affect the strength, number and length distribution of DNA loops compared to Eco1-AID cells in the absence of auxin (Figure 2F and G). Therefore, Eco1 doesn’t appear to play a role in the regulation of DNA-loop sizes after their establishment during S-phase. In other terms, inhibition of cohesin-dependent loops expansion by Eco1 may be restricted to the S phase, shortly after cohesin is being loaded onto DNA during replication, and may no longer occur after S phase completion. It is also possible that the translocase activity required to expand DNA loop is not active during mitosis. Alternatively, Eco1 mediated Smc3 acetylation at S phase may ensure inhibition of DNA loop expansion during the rest of mitosis.

**Figure 2:**
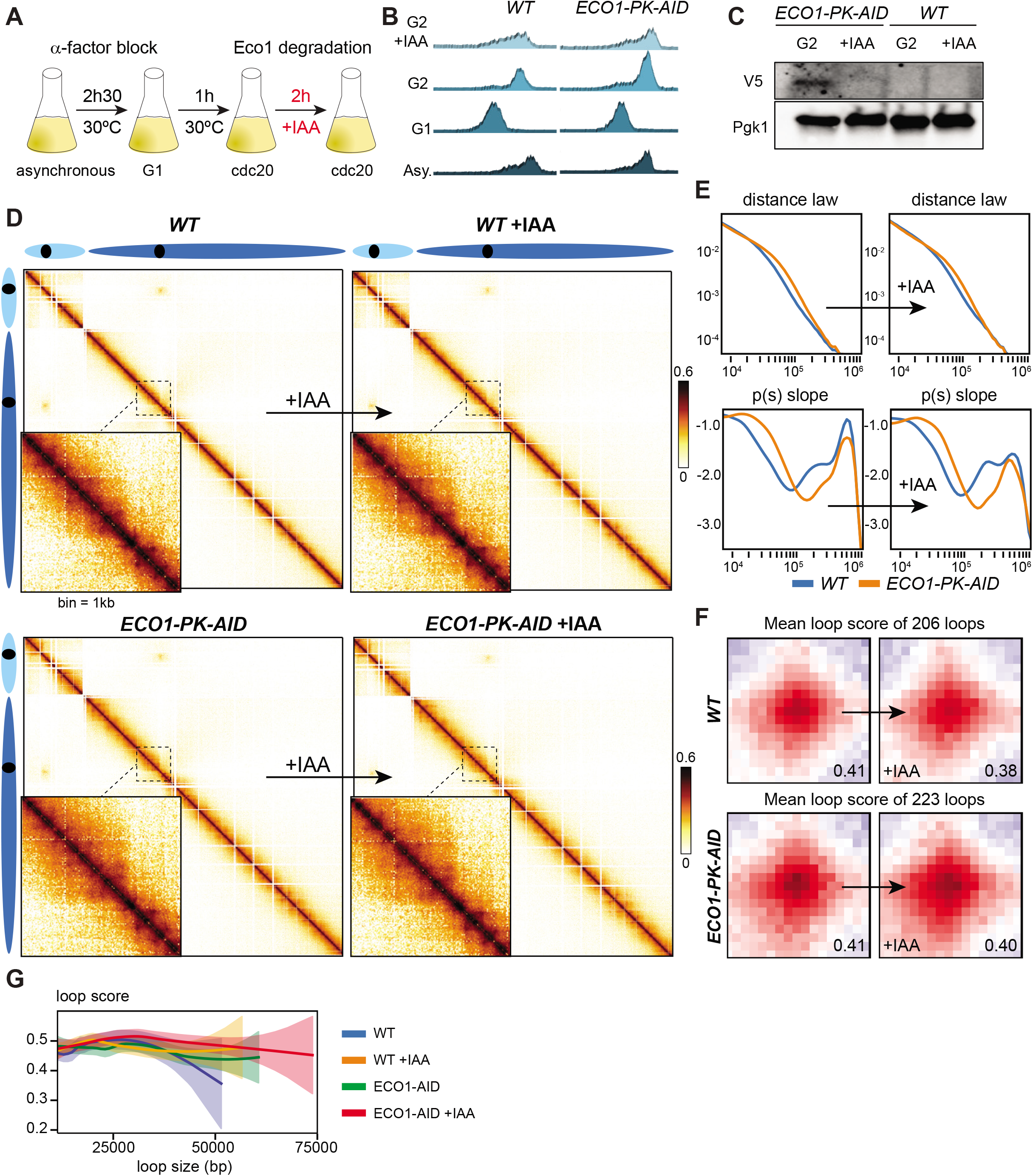
Eco1 is required to restrict DNA loop expansion only during S phase. A. Illustration of the followed experimental protocol to deplete Eco1-AID during metaphase. B. Cell synchronization in G1 and metaphase were confirmed by flow cytometry. C. Western Blot assessing Eco1-pk-AID degradation with anti V5 antibody. Pgk1 was used as loading control. D. Contact maps of chromosomes III and IV (bin 1kb) for metaphase arrested *WT* (top, FB133-57B) and *ECO1-AID* (bottom, FB133-20c) cells before (left) and after (right) auxin (IAA) addition. Chromatin interactions along a region (300kb to 510kb) of the chromosome IV are represented in the black square. E. Contact probability curves Pc(s) representing contact probability as a function of genomic distance (bp) and their respective derivative curves for *WT* and *ECO1-AID* after auxin treatment. F. Mean profile heatmap of loops called by chromosight. G. Loop spectrum indicating scores in function of loop size.

### DNA replication stimulates the expansion of DNA loops

Depletion of the helicase Cdc45, resulting in the progression of unreplicated chromosomes into G2/M, doesn’t prevent chromosome folding, showing that DNA replication is not necessary to regulate the expansion and establishment of DNA loops (Figure 3) (Dauban et al., 2020). However, since replication is essential to trigger Smc3 acetylation, one would have expected that a lack of replication would mimic the effect of Eco1 depletion (i.e. the absence of acetylation), and therefore induce expansion of intrachromosomal contacts to greater distances. To explain this discrepancy, we hypothesized that Eco1 acetylates a sub-population of Smc3 even in the absence of DNA replication. We generated Hi-C maps of mitotic chromosomes from cells that reached mitosis without Eco1 nor Cdc45 (Figure 3A, 3B, 3C, 3D). Contact maps and their associated *p(s*) revealed that Eco1 depletion has only a modest effect on chromatin interactions once DNA replication is repressed (Figure 3D, 3E, 3F, 3G). The effect of Eco1 depletion in the Cdc45 mutant cannot explain on its own the effect which is observed when Eco1 is depleted in replicating cells (Figure 1D, 3E). We propose two hypotheses to explain these observations. On the one hand, active processes favor the extension of DNA loops during DNA replication. On the second hand, active processes to prevent DNA loop expansion are more efficient in Cdc45 depleted cells, i.e. release of cohesin dependent DNA loops mediated by Wpl1.

**Figure 3:**
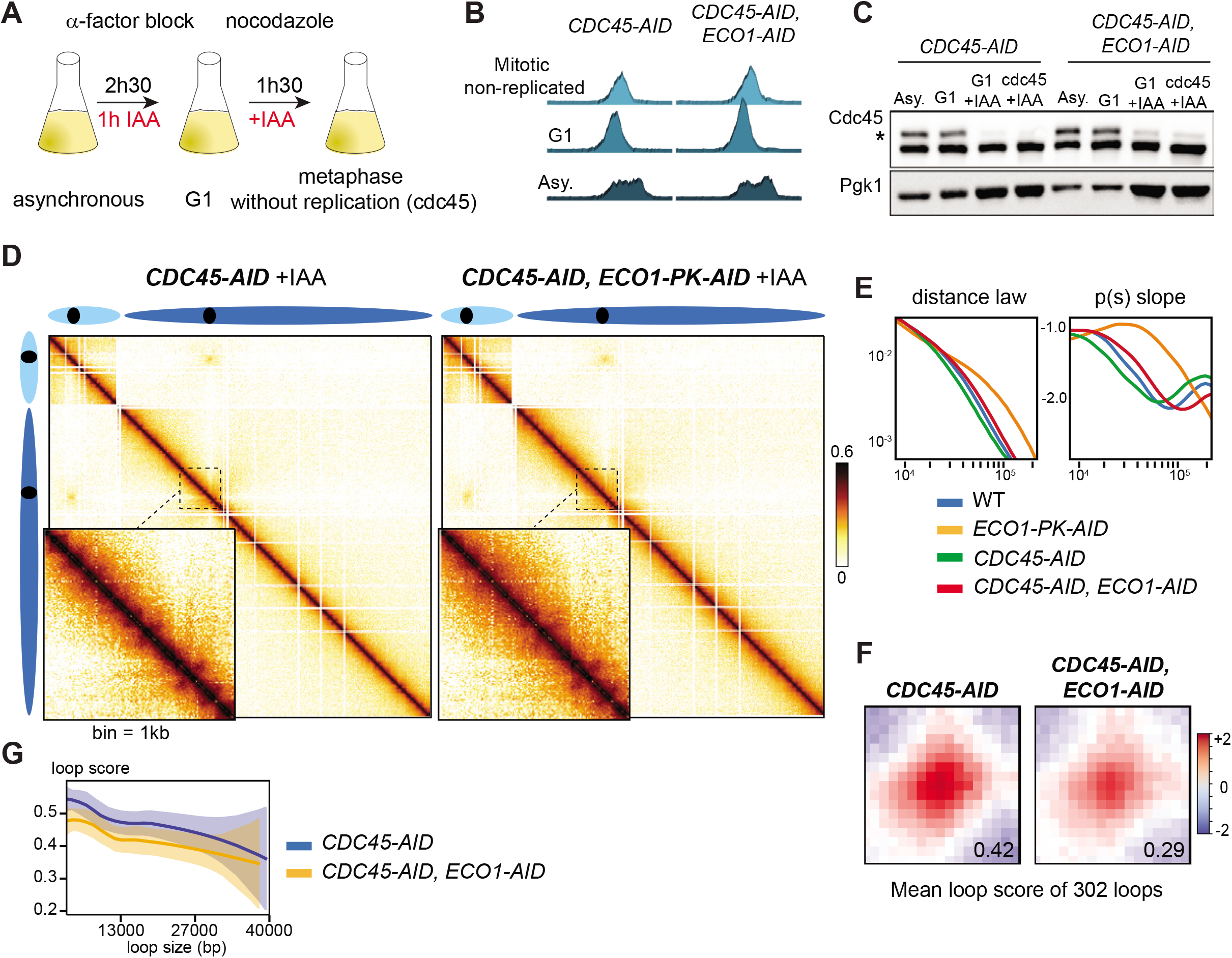
Eco1 has a modest effect in absence of DNA Replication. A. Schematic representation of the protocol followed to degrade Cdc45-AID and Eco1-AID in G1 and arrested cells in G2. B. Cell synchronization was monitored by flow cytometry C. Cdc45 degradation was assessed by western blot using anti-FLAG antibody. Pgk1 was used as loading control. D. Contact maps of chromosomes III and IV (bin 1kb) for metaphase arrested *CDC45-AID* (left, FB154) and *CDC45-AID, ECO1-AID* (right, FB162-1C) cells. E. Intra-chromosomal contact probability as a function of genomic distance and their respective derivative curves for *WT*, (FB133-57B), *ECO1-AID* (FB133-20c), *CDC45-AID* (FB154), and *CDC45-AID, ECO1-AID* (FB162-1C) cells. F. Mean profile heatmap of loops called by chromosight G. Loop spectrum indicating scores in function of loop size

### A subpopulation of DNA loops are stable in the absence of cohesin loading during mitosis

Smc3 acetylation ensures stable cohesin binding on DNA by preventing cohesin removal from DNA by Wpl1 mediated releasing activity. Therefore, if DNA loops on mitotic chromosomes are anchored by acetylated Smc3, they should be resistant to the action of Wpl1 and be maintained in the absence of cohesin *de novo* loading. In order to address the role of the cohesin loader we used *scc2-45*, a thermosensitive allele of Scc2. At permissive temperature, *scc2-45* allows DNA loop formation and chromosome compaction of nocodazole arrested cells (Figure S1A-E, Methods). The number of loops (336 loops), the maximum loop size (45kb) and mean loop score (0.30) are however reduced in comparison to WT in this experimental setup (with 510 loops, 60kb, and MLS= 0.44, respectively; Figure S1F and S1G). Note that the overall number of WT loops detected by Hi-C is increased in the presence of nocodazole (510, Figure S1D and S1F) compared to a cdc20 block (223, Figure 1D): for reasons we don’t fully understand, the contact signal is indeed systematically neater in nocodazole conditions. Nevertheless, and as expected, at non-permissive temperature *scc2-45* fully abolishes cohesin loading, with no DNA loops nor DNA compaction detected while cells replicate their DNA (Figure S1D, S1E and S1F). Contact probabilities (P(s)) following Scc2 inactivation were similar to those of uncondensed G1 cells (Figure S1E) (Lazar-Stefanita et al., 2017), and loop scores of 0.05 or 0.08, similar to G1, were measured after 1h or 2h at restrictive temperature (Figure S1F). In addition, Smc3 acetylation did not take place, as expected from the absence of cohesin loading and subsequent recruitment of Eco1 (Figure S1C). Therefore, at restrictive temperature, *scc2-45* allele fully inactivates the establishment of DNA loops in cycling cells.

To test whether mitotic DNA loops are stable in the absence of *de novo* loading during mitosis, we arrested wild type *SCC2* and *scc2-45* cells in G2/M at permissive temperature (30°C), before shifting both cultures to restrictive temperature (37°C) for either 20 or 60 min (Figure 4A and 4B). This treatment is sufficient to fully inactivate the *scc2-45* allele (Figure S1, see also (Srinivasan et al., 2019). Hi-C maps were generated for both conditions (Figure 4C). Hi-C maps and loop calling analysis revealed that Scc2 inactivation during mitosis has little impact on the positions, strengths (determined by the mean score in Figure 3D) and lengths of DNA loops (Figure 4E). In addition, the number of loops detected in mitosis in these conditions (336 loops) was only modestly affected by 20 min (308 loops) or 60 min (286 loops) of Scc2 inactivation. These results show that a large proportion of mitotic DNA loops are stable in the absence of active Scc2. In addition, while Scc2 inactivation has little effect on DNA loop maintenance, the overall amount of intra-chromosomal contacts, or compaction as seen from the contact probabilities P(s) curves (Figure 4F), and the log ratio between maps (Figure 4G), diminishes.

**Figure 4:**
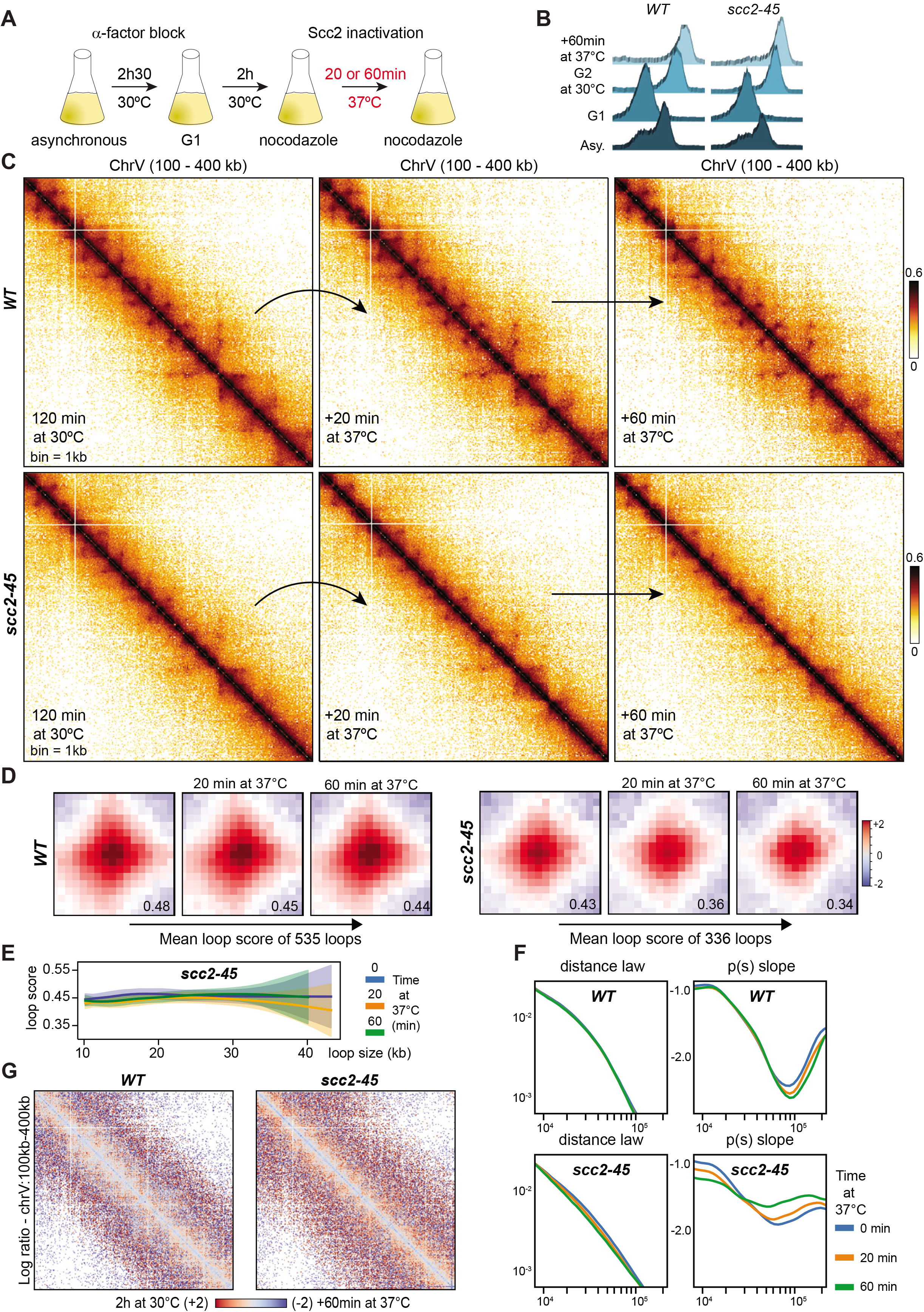
Scc2 is dispensable for maintenance of cohesin dependent loops in G2. A. Schematic representation of the protocol followed to inactivate Scc2 in G2. B. Cell synchronization was monitored by flow cytometry. C. Contact maps of a part of chromosome V (100kb to 400kb) for *WT* (W303-1A) and *scc2-45* (*KN20751*) cells arrested in mitosis at permissive temperature and then shifted at restrictive temperature during 20min or 60min. D. Mean profile heatmap of loops called by chromosight. E. Loop spectrum indicating scores in function of loop size. F. Intra-chromosomal contact probability as a function of genomic distance and their respective derivative curves for *WT* and *scc2-45* cells arrested in mitosis at permissive temperature, and then shifted at 37°C during 20min and 60min. G. Log2 ratio between Hi-C maps from *WT* and *Scc2-45* mitotic cells at permissive or restrictive temperature during 60min.

In other words, while visible, DNA loops on contact maps are maintained after Scc2 inactivation in mitosis, *cis* contacts between loci are reduced. This suggests that during mitosis, the turning over of a non-acetylated form of cohesin may also promote intra-chromosomal contacts in mitosis. We hypothesized that *cis* contacts depending of nonacetylated process may be established during the ongoing of LE process.

### The translocase activity required to expand DNA loop is active during mitosis

We previously observed that depleting Pds5 prior to S phase results in an increase of contacts over longer distances (Dauban et al., 2020). To test whether this increase occurs during mitosis, we depleted Pds5 in post-replicative cells arrested in G2/M by nocodazole using a Pds5-AID degron system (Methods; Figure 5A and 5B). Pds5 degradation was confirmed by western blot (Figure 5C). Hi-C contact maps and P(s) curves revealed that Pds5 depletion results in the spreading of intrachromosomal contacts over longer distances (Figure 5D and 5E) and in the loss of most DNA loops present along yeast mitotic chromosome (411 loops in presence of Pds5 compared to 134 loops in absence of Pds5). The mean loop score was also strongly reduced, from 0,43 to 0,09 (Figure 5F). Not only this result shows that the mechanism driving loop expansion is fully active in post-replicated cells, but it also shows that Pds5, unlike Eco1 (Figure 2), is required to constrain DNA loops during mitosis. The regulation of DNA loops (i.e. sizes and positions) during mitosis, after their establishment, is therefore an active process.

**Figure 5:**
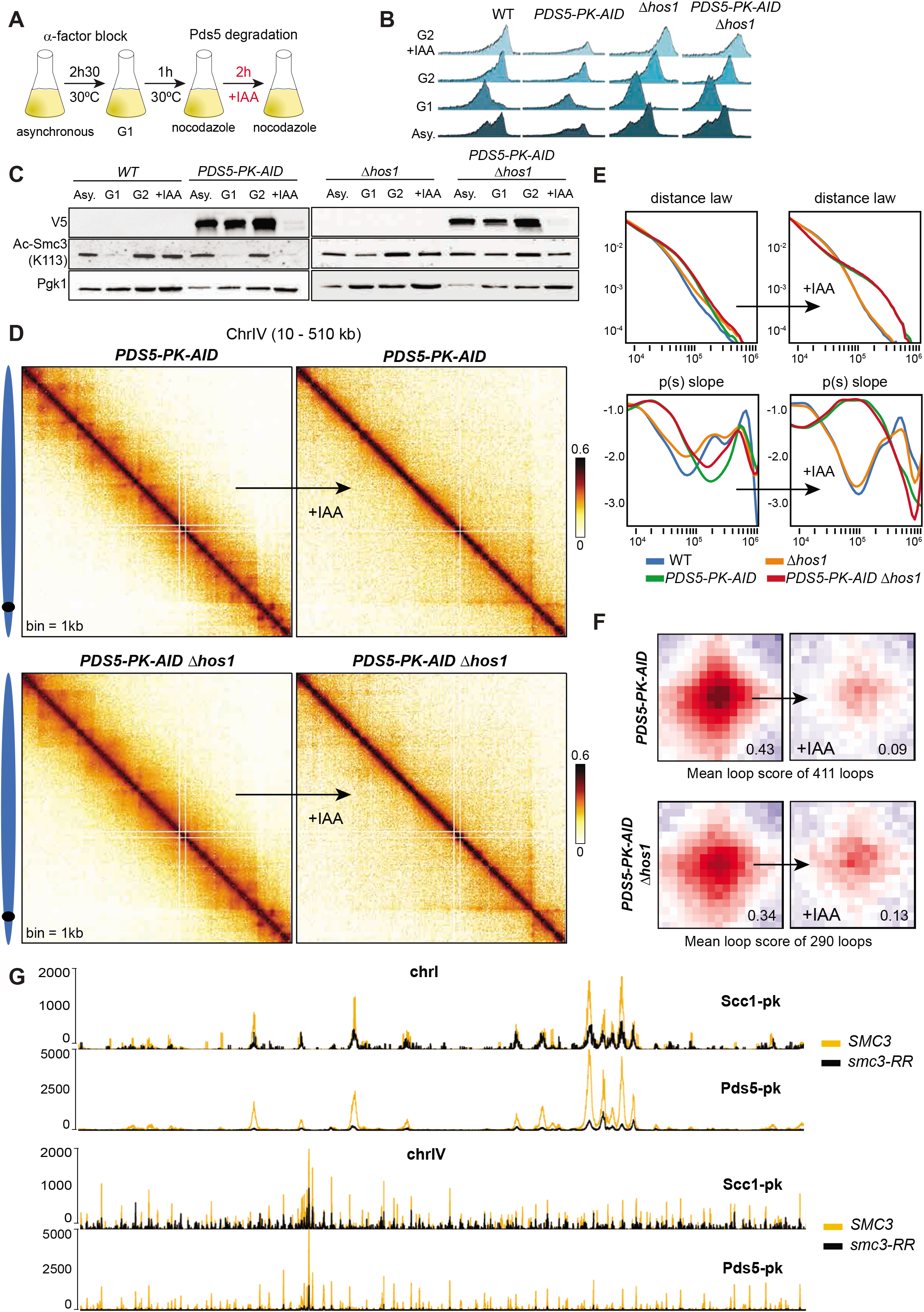
Smc3 acetylation *per se* is not sufficient to block expansion of DNA loop. A. Schema illustrating the experimental protocol for Pds5 degradation in G2. B. Cell synchronisation was monitored by flow cytometry. C. Pds5 degradation and loss of Smc3 acetylation were confirmed by western blot using V5 (Pds5-PK-aid) and Smc3-K113Ac antibodies respectively. Pgk1 was used as loading control. D. Hi-C contact matrices of a part (10kb to 510kb) of chromosome IV (bin = 1kb) showing effect of Pds5 depletion in G2 in presence (yNB33.1-8a) or in absence (yNB40-2b) of Hos1. E. Contact probability curves Pc(s) representing contact probability as a function of genomic distance (bp) and their respective derivative curves. F. Mean profile heatmap of loops called by chromosight. G. Calibrated ChIP-seq profiles showing effect of *smc3-RR* on the distribution of Scc1-PK and Pds5-PK on chromosomes I and IV.

### Scc2 stimulates translocase activity in living cells

We then asked if spreading of intrachromosomal contacts over longer distances observed in Pds5 depleted cells is driven by Scc2. To test this, we used a yeast strain expressing both *scc2-45* and *PDS5-AID* alleles. To address this question, we also introduced in this genetic background a *scc3-K404E* allele that prevents Wpl1-mediated cohesin dissociation from DNA. We noticed that this mutant increases the number of pre-established DNA loops in strain expressing *scc2-45* and *PDS5-AID* alleles and therefore may facilitate the analysis. Pds5 was depleted from cells arrested in G2 and efficient depletion was confirmed by Western blot analysis (Figure 6A, 6B, 6C). Hi-C maps and p(s) curves revealed that spreading of intrachromosomal contacts induced by Pds5 depletion was partially suppressed by the *scc2-45* allele (Figure 6D, 6F). Moreover, according to the MLS, loss of DNA loop was attenuated in *scc2-45* cells (from 0.43 to 0.25) compared to control (from 0,44 to 0,16) (Figure 6E). *Scc2-45* also suppresses the increase of loop size induced by Pds5 depletion (Figure 6G). This data strongly supports the existence of a Scc2 mediated translocation process driving expansion of DNA loops in living cells.

**Figure 6:**
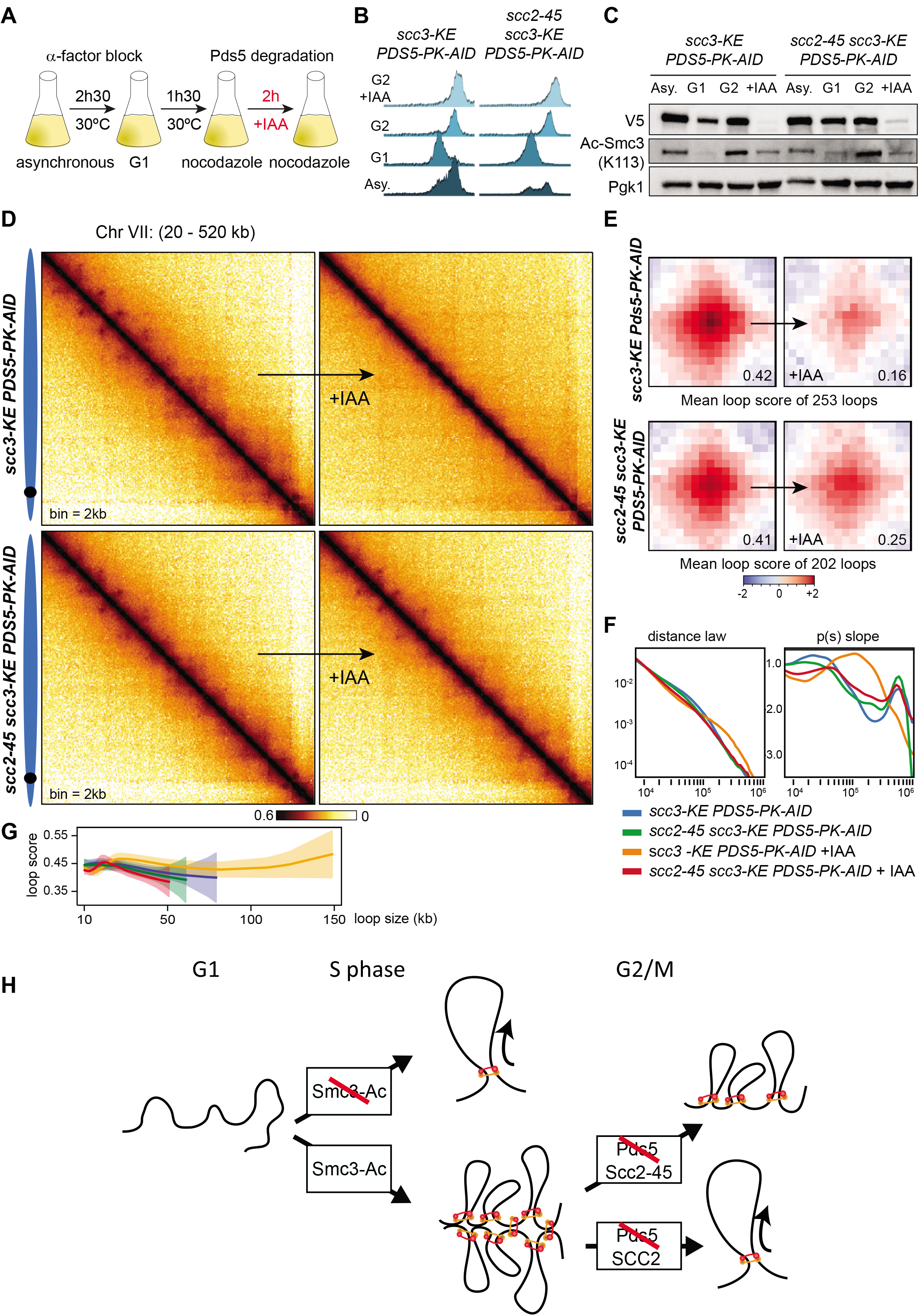
Scc2 drives translocation process. A. Schema illustrating the protocol used to degrade Pds5 in G2. B. Cell synchronisation was monitored by Flow cytometry showing C. Pds5 degradation and loss of Smc3 acetylation in G2 were analysed by western blot V5 (Pds5-PK-aid) and Smc3-K113Ac antibodies respectively. Pgk1 was used as loading control. D. Hi-C contact matrices of a part (20kb to 520kb) of chromosome VII (bin = 2kb) showing effect of Pds5 depletion in *scc3-K404E, scc2-45* (yNB33.2-8c) and *scc3-K404E* (yNB55.1-2a) background. E. Mean profile heatmap of loops called by chromosight. F. Contact probability curves Pc(s) representing contact probability as a function of genomic distance (bp) and their respective derivative curves G. Loop spectrum indicating scores as function of loop size (bp) H. Model

### The block of loop expansion by Smc3 acetylation requires Pds5

Pds5 protects Smc3 from deacetylation by Hos1/HDAC8 in mitosis (Figure 5C) (Chan et al., 2013). To distinguish between the roles of Smc3 acetylation and Pds5 in blocking the expansion of chromatin loops during mitosis, we asked whether Smc3 acetylation inhibits loop expansion in a Pds5 dependent or independent mechanism. To test this, we prevented Smc3 deacetylation in a *hos1Δ* background prior degrading Pds5 in mitosis (Figure 5B and 5C). In other words, we tested whether preventing Smc3 deacetylation would stop, or slow down, the DNA loop expansion resulting from Pds5 depletion in G2/M arrested cells. We note that hos1 deletion had no significant impact on DNA loop formation (Figure S2B and S2C). Comparison of Hi-C contact maps, revealed that the absence of Hos1/HDAC8 does not suppress the effect of Pds5 depletion on chromatin interaction (Figure 5D), as confirmed by P(s) curves and DNA loop mean score (Figure 5E and 5F). Our results therefore suggest that Smc3 acetylation is not sufficient *per se* to block DNA loop expansion independently of Pds5.

### Smc3 acetylation stabilizes Pds5 on DNA at loop borders

As Eco1 activity is thought to stabilize the cohesin subunit Pds5 on DNA, we asked whether Smc3 lysines K112 and K113 are the targets of Eco1 that improve Pds5 binding on cohesin and consequently anchor DNA loops. To check this hypothesis, we measured the effect of non-acetylable Smc3 on Pds5 versus Scc1’s association with the entire genome using calibrated ChIP-seq. To avoid any confusing variables due to Wpl1-mediated releasing activity, all ChIP-seq experiments were performed in a cellular context depleted for Wpl1. Scc1 peak heights observed in *Smc3-RR* condition were slightly reduced compared to Smc3 WT as peaks were also broadened (Figure 5G and S2C). This cohesin distribution may be a consequence of the translocation process that passes the former “stop sites” in *smc3-RR* cells (Figure 5G and S2D). In addition, Pds5 binding was highly reduced in *smc3-RR* condition compared to WT (Figure 5G and S2D) showing that Smc3 K112 and K113 are essential for Pds5 positioning at cohesin sites. Occupancy ratios were calculated as in (Hu et al., 2015) and were 1,609 in *SMC3 WT* and 1,399 in *smc3-RR* for Scc1-pk. They were 1,087 in *SMC3 WT* and 0,167 in *smc3-RR* for Pds5-pk (Figure S2D). Therefore, these results support the notion that Smc3 acetylation stabilizes Pds5 at DNA loop anchors.

## DISCUSSION

### Eco1-dependent acetylation of Smc3 inhibits loop expansion

Recent studies have brought forward the idea that cohesin organizes mammalian interphase and yeast metaphase chromosomes by extruding DNA loops, a process which consists in capturing chromatin loops within the cohesin ring and progressively enlarging them up to Mb-sized structures. The molecular factors that regulate cohesin mediated loop expansion are related to those involved in the establishment and maintenance of SCC. Experiments on mammalian and yeast cells showing that chromatin loop lengths increase when Wpl1 is inactivated have suggested that its releasing activity restricts DNA loop expansion by removing cohesin from the DNA (Dauban et al., 2020; Haarhuis et al., 2017). Moreover, we and others demonstrated that an Eco1-dependent mechanism also inhibits DNA loop expansion in both human and yeast (Dauban et al., 2020; Wutz et al., 2020). Inactivation of Eco1 also induces the enlargement of loops and spreading of intrachromosomal contact over greater distances. According to the loop extrusion model, contacts over longer distances can result either from an increase of cohesin residence time on the DNA, or from a stimulation (or de-repression) of a translocase activity. As the absence of Eco1 reduces the cohesin residence time (Chan et al., 2012), we postulated that the long range, intrachromosomal cis contacts observed in Eco1 depleted cells may rather arose from a stimulation, or de-repression, of a translocase activity. In the present work we show that preventing the acetylation of K112 and K113, two conserved lysine within Smc3 ATPase head (*Smc3-RR* mutant), also lead to longer cis contacts, a phenotype similar to that of Eco1 mutant. During replication, these two Smc3 lysines are acetylated by Eco1 to ensure SCC (Chan et al., 2012; Rolef Ben-Shahar et al., 2008; Rowland et al., 2009; Unal et al., 2008; Zhang et al., 2008). Finally, our results show that Eco1 depletion has little effect in *cdc45* cells that are G2/M arrested without going through replication. This strongly suggest that Eco1-dependent acetylation of Smc3 during S phase inhibits the translocation processes (Figure 6H).

### How does cohesin expand DNA loops?

To understand how Smc3 acetylation limits the length of DNA loops *in vivo*, it is crucial to decipher the mechanisms that stimulate the expansion of these loops. The inactivation of Pds5 in G2, after replication, induces the expansion of pre-established loops (Figure 5). It has been proposed that stimulation of cohesin ATP hydrolysis by Scc2 (NIPBL in mammals) promotes the translocase activity supporting DNA loop expansion. Indeed, depleting NIPBL led to TADs disruption and decreased DNA loop size in either primary or haploid mammalian cells (Haarhuis et al., 2017; Schwarzer et al., 2017). In both systems however, it was impossible to distinguish a role for Scc2 as a potential cohesin translocase from its conventional role in cohesin loading, that would also affect the residence time parameter and disturb the loop patterns. The importance of Scc2 in promoting DNA loop extrusion also comes from in vitro data (Davidson et al., 2019; Gutierrez-Escribano et al., 2019). Crucially, we show that Scc2 promotes DNA loop expansion *in vivo* in the absence of Pds5, after cohesin are loaded on DNA. The *scc2-45* allele abolishes this stimulation showing that Scc2 is required for the translocation process (Figure 6H).

### Is Scc2 required to maintain DNA loops?

Our present work reveals that a large amount of pre-formed DNA loop is preserved after Scc2 inactivation in G2 arrested cells, showing that most loops are maintained in absence of cohesin loading. However, the length of a significant portion of intrachromosomal contacts is reduced following Scc2 inactivation as chromosomes decondense (Figure 4). This indicates that Scc2 is needed to maintain a sub-population of *cis* chromosomal contacts along the genome. We postulate that during mitosis, Scc2 may be required to *de novo* load nonacetylated cohesin complexes to promote intra-chromosomal contacts. It can also be envisaged that Scc2 is necessary to maintain a portion of cohesin engaged in cis contacts on DNA. It is possible that Scc2 renders cohesin resistant to releasing activity by avoiding binding of Pds5/Wpl1 on the kleisin subunit or counteracts a Wpl1 independent pathway (Srinivasan et al., 2019). Recent structural studies also revealed that engagement of ATPase head prior to ATPase hydrolysis dissociates Scc1 from Smc3 (Higashi et al., 2020). Opening of this interface during translocation process may induce escape of DNA from cohesin. Therefore, Scc2 may avoid release of cohesin from DNA by preventing opening of Scc1-Smc3. Additional investigations are necessary to fully understand all Scc2 dependent mechanisms regulating LE process.

### How does Smc3 acetylation inhibits loop expansion?

Structural studies from fission yeast and human revealed that NIPBL/Scc2 interacts with the conserved acetyl acceptor lysine of Smc3 to trigger ATP hydrolysis (Higashi et al., 2020; Shi et al., 2020). It has been proposed that ATP hydrolysis stimulated by Scc2 promotes the translocase activity supporting DNA loop expansion, and that acetylation of Smc3 K112 and K113 *per se* may compromise the translocation process. This idea was based on the facts that changing KK to QQ reduces stimulation of ATPase activity by Scc2 *in vitro* (Collier et al., 2020). Based on structural data it was proposed that Smc3 acetylation abolishes Scc2 binding to cohesin. However, our data shows that Smc3 acetylation isn’t sufficient to block DNA loop expansion in the absence of Pds5, indicating that Smc3 acetylation *per se* is not sufficient to block Scc2 mediated ATPase hydrolyses.

Pds5 is thought to compete with Scc2 for the binding on cohesin (Kikuchi et al., 2016; Petela et al., 2018) Pds5 and Scc2 have a similar overall structure, suggesting that they may bind to the same regions on the cohesin ring (Higashi et al., 2020; Lee et al., 2016; Muir et al., 2016; Ouyang et al., 2016; Shi et al., 2020). Eco1 inactivation also affects the way that Pds5 binds to chromatin (Chan et al., 2013; Chapard et al., 2019). Crucially we showed that the amount of Pds5 on chromatin decreases in cells expressing Smc3-RR instead of Smc3-KK. This indicates that Pds5 may also interact with acetyl acceptor lysines of Smc3 or close to their location. Smc3 acetylation of those lysines may strengthen Pds5 binding to cohesin, and consequently inhibits Scc2 mediated ATPase hydrolysis.

In addition of leading to contacts over longer distances, *smc3-RR* reduces the number of DNA loops at discrete positions. This indicates that Smc3 acetylation also positively regulates the maintenance of cohesin-dependent DNA loops. Therefore, by acetylating Smc3 on lysine residues K112 and K113, Eco1 stabilises cohesin units on DNA and not only entitles SCC, but also maintains DNA loops at discrete positions. However, although the underlying mechanisms appear similar, relying in part on Smc3 acetylation, they remain distinguishable as the latter can occur in the absence of the former. We postulate that acetylation of cohesin extruding DNA loops helps anchoring and maintaining DNA loops at discrete positions along the genome. Nevertheless, LE process could also be stopped by a portion of cohesin which is acetylated and that topologically entraps one or two sister DNA. In other terms, stably bound cohesin involved in SCC may also halt extrusion process and consequently induce appearance of additional DNA loops along the genome. The factors that regulate DNA loop stability in cdc45 cells have yet to be explored.

Our present findings suggest that Smc3 acetylation is needed to stabilize loops at discrete positions by reinforcing Pds5 binding onto cohesin bound DNA (Figure 6H). Our data showing that Pds5 depletion decreases the number of DNA loops also supports this notion. Pds5 may create a compartment in which DNA is being trapped, similarly to what is observed for Ycg1 subunit that anchors condensin oligomers on DNA (Kschonsak et al., 2017). This anchoring by Pds5 is compatible with the role of Pds5 in providing and maintaining cohesion (Chan et al., 2013) but also with either one sided or double-sided DNA loop extrusion (Banigan et al., 2020).

## Acknowledgments

This research was supported by the European Research Council under the Horizon 2020 Program (ERC grant agreement 771813 to RK), the Fondation ARC pour la Recherche sur le Cancer (ARCPJA22020060002067 to FB) and the Comité de l’Occitanie de la Ligue Nationale contre le Cancer (to FB). Nathalie Bastié and LD were supported by the Ministère de l’Enseignement Supérieur et de la Recherche. Christophe Chapard was supported by Pasteur-Roux-Cantarini fellowship.

We thank all our colleagues from the laboratories regulation spatiale des génomes and organization du noyau, especially Cyril Matthey-Doret. We also thank Martin Houlard, Jean Paul Javerzat for helpful comments on the manuscript, Kim Nasmyth’s laboratory for strains and K. Shirahige for the Smc3 antibody.

## Author contributions

NB, CC, FB and RK designed research. NB and CC performed the experiments, with contributions from LD for the early stages of the project. CC analyzed the data. All authors interpreted the data. NB, CC, RK, FB wrote the manuscript.

## Data Availability

Sample description and raw sequences are accessible on SRA database through the following accession number: PRJNA715343

## SUPPLEMENTARY DATA

**Figure S1:**
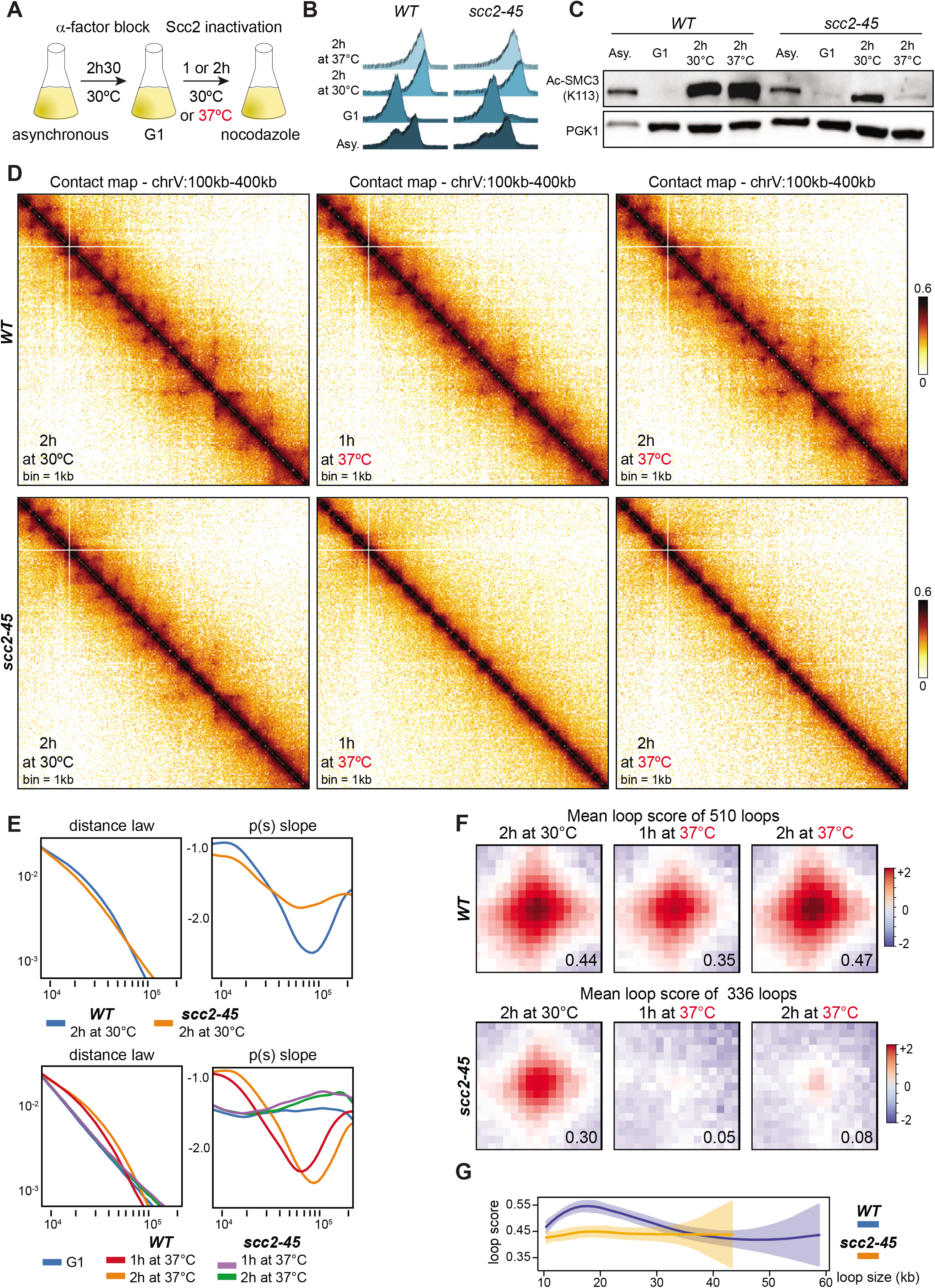
*scc2-45* allele fully inactivates establishment of DNA loop (related to Fig. 4) A. Schematic representation of the protocol followed to inactivate Scc2 during S phase. B. Cell synchronization was monitored by flow cytometry. C. Western blotting showing that inactivation of Scc2 prior to S phase induces loss of Smc3 acetylation D. Contact maps of a part of chromosome V (100kb to 400kb) for *WT* (W303-1A) and *scc2-45* (KN20751) cells arrested in mitosis at permissive temperature during 2h or arrested at restrictive temperature during 1h or 2h. E. Intra-chromosomal contact probability as a function of genomic distance and their respective derivative curves for *WT* and *scc2-45* cells arrested in mitosis at permissive temperature during 2h or arrested at restrictive temperature during 1h or 2h F. Mean profile heatmap of loops called by chromosight. G. Loop spectrum indicating scores in function of loop size.

**Figure S2:**
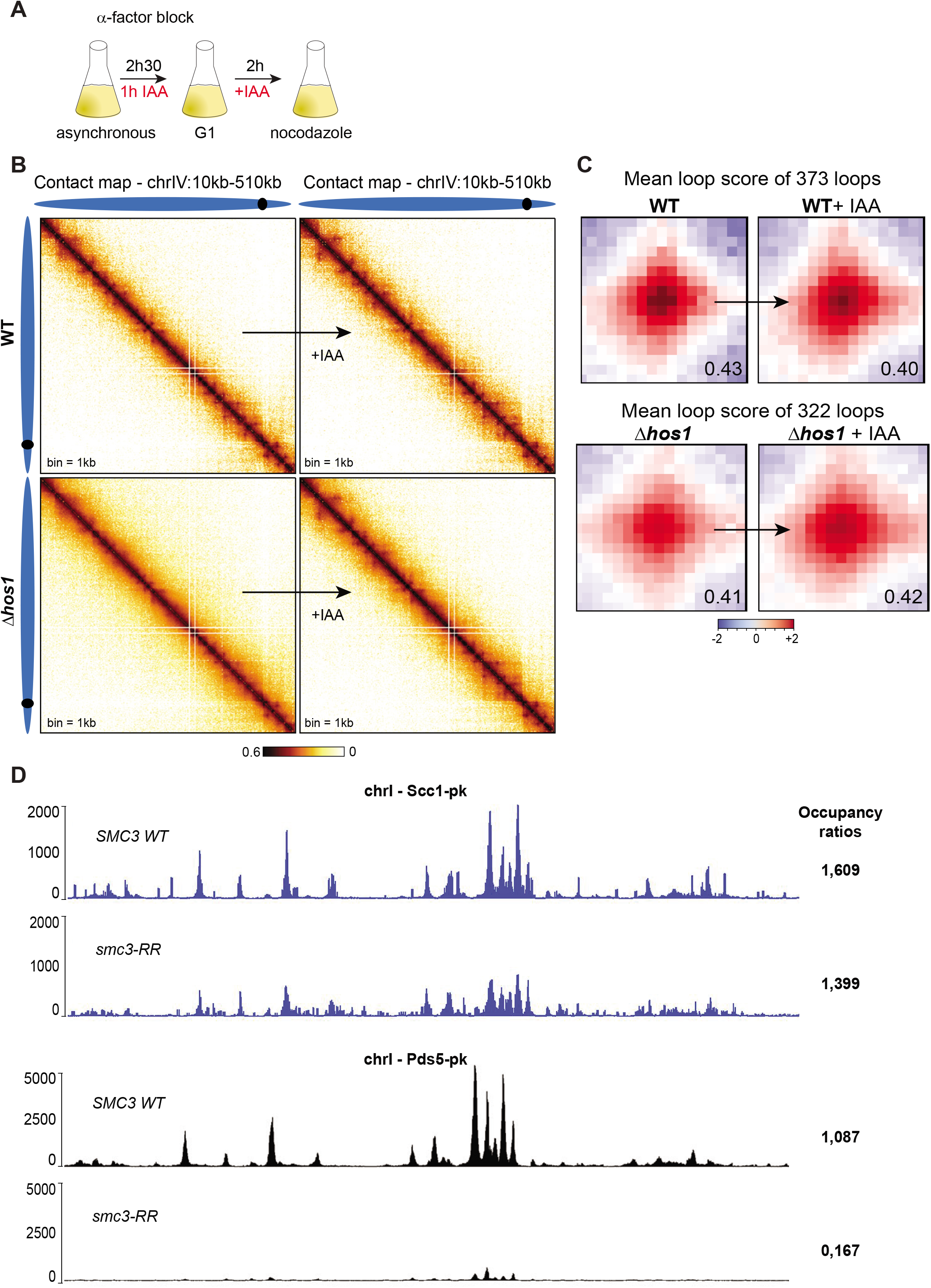
Effect of Hos1 inactivation on chromatin interactions (related to Fig. 5) A. Schematic representation of the protocol followed to synchronize cells. B. Hi-C contact matrices of a part (10kb to 510kb) of chromosome IV (bin = 1kb) of WT (W303-1A) and *hos1Δ* (yNB40-1c) cells. C. Mean profile heatmap of loops called by chromosight. D. Calibrated ChIP seq profiles showing effect of *smc3-RR* on the distribution of Scc1-PK and Pds5-PK on chromosome I and the respective occupancy ratios

## EXPERIMENTAL PROCEDURES

### Media culture conditions and synchronisation

All strains used in this study are derivate of W303 and are listed in the Table “Strain list”. All strains were grown overnight at 30°C or 25°C in 150mL of suitable media to attain 4,2×10^8 cells.

Degradation of ECO1-AID and SMC3-AID in G1 (Figure 1)

To study effect of Eco1 inactivation and *smc3-RR* on chromosome organisation, overnight culture of the strains, FB133-20C (*ECO1-AID*), yNB54-4 (*SMC3, SMC3-AID*), yNB54-3 (*smc3-RR. SMC3-AID*), FB134-16C (*SMC3-AID*) and yLD116-1a (*cdc20*) were gown in a complete medium deprived of methionine (SC-met) (SC: 0.67% yeast nitrogen base without amino acids (Difco)), supplemented with a mix of amino-acids, uracil and adenine, 2% glucose). To arrest cells in metaphase, yeast cells were synchronised in G1 by adding of α-factor (Antibodies-online, ABIN399114) in the media every 30min during 2h30 (1μg/mL final). Auxin (2mM final) (Sigma-Aldrich, I3750) was added to the media 1h after starting α-factor arrest. After G1 arrest cells were washed twice in fresh YP and released in rich medium (YPD): 1%bacto pepone (Difco), 1% bacto yeast extract (Difco), and 2% glucose) supplemented with 2mM of methionine and containing auxin.

Degradation of ECO1-AID in G2 (Figure 2)

To analyse effect of Eco1 depletion in metaphase on chromatin interactions strains FB133-57B (*cdc20*) and FB133-20C (*ECO1-AID*) were grown overnight in a complete medium deprived of methionine and then arrested in G1 using α-factor. After 2h30 in G1, cells were washed twice in fresh YP and released in rich medium (YPD) supplemented in methionine (2mM final) to be re-arrested in metaphase. After 2 h in metaphase, 2mM of auxin was added during two more hours.

Depletion of CDC45-AID and ECO1-AID (Figure 3)

To study effect of DNA replication on chromatin loops expansion, strains *FB154(CDC45-AID*) and FB162-1C (*CDC45-AID. ECO1-AID*), were grown overnight in YPD. Cells were synchronised in G1 using α-factor during 2h30, auxin (2mM final) was added to the media 1h after starting α-factor arrest. After G1 arrest, cells were washed in fresh YP and released in rich medium (YPD) containing Nocodazole (Sigma-Aldrich, M1404-10MG) and auxin (2mM final) during 1h30.

Inactivation of Scc2 (Figure 4 and S1)

To check effect of Scc2 inactivation on establishment of cohesin-dependent loops cells KN20751(*scc2-45*) and W303-1A (*WT*) were synchronised in G1 thanks to α-factor at 30°C during 2h30. Then yeasts cells were washed and released in YPD containing Nocodazole at 37°C during 1h or 2h. To determine effect of Scc2 inactivation on maintenance of cohesin dependant loop, KN20751 (*scc2-45*) and W303-1A (*WT*) was first arrest in G1 thanks to α-factor as described previously, then cells were released in G2 with Nocodazole at 30°C during 2h. After G2 arrest, cells in media containing Nocodazole were heat in a water bath at 37°C during 20min or 60min.

Degradation of Pds5 (Figure 5 and S2)

To analyse effect of Pds5 depletion in G2, strains yNB33.1-8a (*PDS5-AID)*, W303-1A (*WT)*, yNB40-2B (*Hos1Δ, PDS5-AID*) and yNB40-1C (*Hos1Δ*) were grown overnight in YPD. Cells were then synchronised in G1 using α-factor and released in YPD media containing Nocodazole to arrest cells in G2 during 1h. After G2 arrest, 2mM of auxin are added during 2h to induce Pds5 degradation.

Depletion of Pds5 in Scc2-45 background (Figure 6)

To study effect of Scc2 on translocase activity, strains yNB33.2-8c (*scc2-45, PDS5-AID, scc3-K404E*) and yNB55.1-2a (*PDS5-AID, scc3-K404E*) were grown overnight in YPD. Cells were synchronised using α-factor during 2h30 and released in YPD containing Nocodazole during 1h30. After G2 arrest, 4mM of auxin was added to the media during 2h to induced Pds5 degradation.

After synchronization all strains were fixed for Hi-C experiments.

### Flow cytometry

To verify cell cycle arrest and synchronisation, 1mL of cells culture were fixed in ethanol 70% and stored overnight at −20°C. Pellet was incubated with 50mM Tris-HCl (pH 7,5) and 5μL RNase A (10mg/mL) overnight at 37°C. Cells were pelleted and resuspended in 400μL of FACS buffer (1mg/mL propidium iodide (Fisher, P3566), Tris-HCl, NaCl, MgCl2) and incubated at 4°C. Cells were sonicated with 60% output for 10secondes.

Flow cytometry was performed on a CyFlow ML Analyzer (Partec) and data were analysed using FloMax software with measure 1000 events at 300events/sec.

### Proteins extractions and acetylation assays

A pellet form 10mL of culture was frozen in liquid nitrogen and stored overnight at −20°C. Cells pellet were resuspended in 100μL H2O and 20μL trichloroacetic acid (Sigma-Aldrich, T8657) (TCA). Then cells were broken by glass beads at 4°C and precipitated proteins were resuspended in Lamely buffer with 100mM DTT and Tris-HCl (pH 9,5). Proteins were extracted by cycles of 5 min heating at 80°C and 5min vortexing at 4°C. After centrifugation, extracted proteins were collected and freeze at −20°C.

Eluates were analysed by SDS-page followed by wester blotting with antibodies anti Smc3-K113Ac (Beckouët et al., 2010), anti V5-tag (VWR, MEDMMM-) and Anti-pgk1(Invitrogen, 459250)

### Calibrated ChIP-sequencing

Cells were grown exponentially to OD600 = 0.5. In triplicates, 15 OD600 units of *S. cerevisiae* cells were mixed with 3 OD600 units of *C. glabrata* to a total volume of 45 mL and fixed with 4mL of fixative solution (50 mM Tris-HCl, pH 8.0; 100 mM NaCl; 0.5 mM EGTA; 1 mM EDTA; 30% (v/v) formaldehyde) for 30 min at room temperature (RT) with rotation. The fixative was quenched with 2mL of 2.5M glycine (RT, 5 min with rotation). The cells were then harvested by centrifugation at 3,500 rpm for 3 min and washed with ice-cold PBS. The cells were then resuspended in 300 mL of ChIP lysis buffer (50 mM HEPES KOH, pH 8.0; 140 mM NaCl; 1 mM EDTA; 1% (v/v) Triton X-100; 0.1% (w/v) sodium deoxycholate; 1 mM PMSF; 2X Complete protease inhibitor cocktail (Roche)) and transfer in tubes 2mL containing glass beads before mechanical cells lysis. The soluble fraction was isolated by centrifugation at 2,000 rpm for 3min then transferred to sonication tubes and samples were sonicated to produce sheared chromatin with a size range of 200-1,000bp. After sonication the samples were centrifuged at 13,200 rpm at 4°C for 20min and the supernatant was transferred into 700μL of ChIP lysis buffer. 80μL (27μl of each sample) of the supernatant was removed (termed ‘whole cell extract [WCE] sample’) and store at −80°C. 5mg of antibody (anti-PK) was added to the remaining supernatant which is then incubated overnight at 4°C (wheel cold room). 50μL of protein G Dynabeads was then added and incubated at 4°C for 2h. Beads were washed 2 times with ChIP lysis buffer, 3 times with high salt ChIP lysis buffer (50mMHEPES-KOH, pH 8.0; 500 mM NaCl; 1 mM EDTA; 1% (v/v) Triton X-100; 0.1% (w/v) sodium deoxycholate;1 mM PMSF), 2 times with ChIP wash buffer (10 mM Tris-HCl, pH 8.0; 0.25MLiCl; 0.5% NP-40; 0.5% sodium deoxycholate; 1mM EDTA;1 mMPMSF) and 1 time with TE pH7.5. The immunoprecipitated chromatin was then eluted by incubation in 120μL TES buffer (50 mMTris-HCl, pH 8.0; 10 mM EDTA; 1% SDS) for 15min at 65°C and the supernatant is collected termed ‘IP sample’. The WCE samples were mixed with 40μL of TES3 buffer (50 mM Tris-HCl, pH 8.0; 10 mM EDTA; 3% SDS). ALL (IP and WCE) samples were de-cross-linked by incubation at 65°C overnight. RNA was degraded by incubation with 2μL RNase A (10 mg/mL) for 1h at 37°C. Proteins were removed by incubation with 10μL of proteinase K (18 mg/mL) for 2h at 65°C. DNA was purified by a phenol/Chloroform extraction. The triplicate IP samples were mixed in 1 tube and libraries for IP and WCE samples were prepared using Invitrogen TM Collibri TM PS DNA Library Prep Kit for Illumina and following manufacturer instructions. Paired-end sequencing on an Illumina NextSeq500 (2×35 bp) was performed.

### Hi-C procedure and sequencing

Cell fixation with 3% formaldehyde (Sigma-Aldrich, Cat. F8775) was performed as described in Dauban et al. Quenching of formaldehyde with 300 mM glycine was performed at room temperature for 20 min. Hi-C experiments were performed with a Hi-C kit (Arima Genomics) with a double DpnII + HinfI restriction digestion following manufacturer instructions. Samples were sonicated using Covaris (DNA 300bp). Preparation of the samples for paired-end sequencing on an Illumina NextSeq500 (2×35 bp) was performed using Invitrogen TM Collibri TM PS DNA Library Prep Kit for Illumina and following manufacturer instructions.

### Processing of the reads, computation of contact matrices, and generation of contact maps

Reads were aligned and the contact data processed using Hicstuff, available on Github (https://github.com/koszullab/hicstuff). Briefly, pairs of reads were aligned iteratively and independently using Bowtie2 in its most sensitive mode against the S. cerevisiae S288C reference genome. Each uniquely mapped read was assigned to a restriction fragment. Quantification of pairwise contacts between restriction fragments was performed with default parameters: uncuts, loops and circularization events were filtered as described in ref. (Cournac et al., 2012) PCR duplicates (defined as paired reads mapping at exactly the same position) were discarded. Contact maps were generated with the “view” function of Hicstuff and normalized using the balance function of Cooler. Bins were set at 1, exp0.2 transformed, and rendered.

### Generation of the ratio maps

Ratio maps were generated with the “view” function of Hicstuff

### Computation of the contact probability as a function of genomic distance

Computation of the contact probability as a function of genomic distance P(s) and its derivative have been determined using the “distance law” function of Hicstuff with default parameters, averaging the contact data of entire chromosome arms.

### Loop detection and scoring with Chromosight

Chromosight 1.3.1 (Matthey-Doret et al., 2020) was used to call loops de novo from contact maps binned at 1 kb and balanced with Cooler (Abdennur and Mirny, 2020). Matrices were subsampled to contain the same total number of contacts. De novo loop calling was computed using the “ detect” mode of Chromosight, with minimum loop length set at 2kb, percentage undetected set at 25 and pearson correlation threshold set at 0.315.

Loop strength was quantified for each loop using the quantify mode of Chromosight and the mean loop score was calculated for each condition. Loop pile-up of averaged 17kb windows were generated with Chromosight.

### Data and software accessibility

The accession number for the sequencing reads reported in this study is PRJNA715343

Open-access versions of the programs and pipeline used (Hicstuff, Chromosight) are available online on the github account of the Koszul lab (https://github.com/koszullab/).

Bowtie2 2.3.4.1 is available online at http://bowtie-bio.sourceforge.net/bowtie2/

STRAINS used in this study

| Yeast strain name | Genetic background | Genotype | Reference |
| --- | --- | --- | --- |
| FB134-16c | W303 | MATa, his3::ADH1promoter-OsTIR1-9myc::HIS3, SMC3-3sIAA::ADE2,TRP1:Metp:CDC20 | Dauban et al. 2020 |
| yNB54-3 | W303 | MATa, his3::ADH1promoter-OsTIR1-9myc::HIS3, leu2::smc3 K112R,K113R::LEU2, SMC3-3sIAA::Kan,TRP1:Metp:CDC20 | This study |
| yNB54-4 | W303 | MATa, his3::ADH1promoter-OsTIR1-9myc::HIS3,SMC3-3sIAA::Kan, Leu2::SMC3::LEU2, TRP1:Metp:CDC20 | This study |
| yLD116-1a | W303 | MATa , TRP1:Metp:CDC20 | This study |
| FB133-20c | W303 | MATa, ura::ADH1 promoter-OsTIR1-9myc::URA3, Eco1-PK3-aid::KanMx4, TRP1:Metp:CDC20 | Dauban et al. 2020 |
| FB133-57B | W303 | MATa, ura::ADH1 promoter-OsTIR1-9myc::URA3, TRP1:Metp:CDC20 | Dauban et al.2020 |
| FB154 | W303 | MATa, CDC45-Flagx5-aid::KanMX4,his3::ADH1promoter-OsTIR1-9myc::HIS3 | Dauban et al.2020 |
| FB162-1C | W303 | MATa, CDC45-Flagx5-aid::KanMX4, ura::ADH1 promoter-OsTIR1-9myc::URA3, Eco1-PK3-aid::KanMx4 | This study |
| W303-1A | W303 | MATa | Rodney Rothstein |
| KN20751 | W303 | MATa, scc2-45::natMX (L545P D575G) | Petela et al. 2018 |
| yNB33.1-8a | W303 | MATa, ura::ADH1promoter-OOs-Tir1-9myc::URA3, Pds5-pk-aid::KanMX4 | This study |
| yNB40-2B | W303 | MATa, hos1D::KanMx:ADE, ura::ADH1promoter-OOs-Tir1-9myc::URA3, Pds5-pk-aid::KanMX4 | This study |
| yNB33.2-8c | W303 | MATa, ura::ADH1 promoter-OOs-TIR1-9myc::URA3, PDS5-Pk3-aid::KanMX4, scc2-45::natMX (L545P D575G), SCC3(K404E)::HA::HIS | This study |
| yNB55.1-2a | W303 | MATa, ura::ADH1 promoter-OOs-TIR1-9myc::URA3, PDS5-Pk3-aid::KanMX4, SCC3(K404E)::HA::HIS | This study |
| yNB40-1C | W303 | MATa, hos1D::KanMx:ADE, ura::ADH1promoter-OOs-Tir1-9myc::URA3 | This study |
| yNB66.1-6b | W303 | MATa, his3::ADH1promoter-OsTIR1-9myc::HIS3, SMC3-3sIAA::Kan, Leu2::SMC3::LEU2, Pds5-PK6::KanMX, Rad61 AID::natMX | This study |
| yNB67.1-1b | W303 | MATa, his3::ADH1promoter-OsTIR1-9myc::HIS3, SMC3-3sIAA::Kan, leu2::smc3K112R,K113R::LEU2, Pds5-PK6::KanMX4, Rad61-AID::natMX | This study |
| yNB64.1-17b | W303 | MATa, his3::ADH1promoter-OsTIR1- | This study |
|  |  | 9myc::HIS3,SMC3-3sIAA::Kan,<br>Leu2::SMC3::LEU2, Scc1-PK3::KanMX, Rad61-AID ::natMX |  |
| yNB65 | W303 | MATa, his3::ADH1promoter-OsTIR1-9myc::HIS3,SMC3-3sIAA::Kan,<br>leu2::smc3 K112R,K113R::LEU2,<br>Scc1-Pk ::KanMX, Rad61-AID::natMX | This study |

## REFERENCES

Abdennur, N., and Mirny, L.A. (2020). Cooler: scalable storage for Hi-C data and other genomically labeled arrays. Bioinformatics 36, 311–316.

Arnould, C., Rocher, V., Finoux, A.L., Clouaire, T., Li, K., Zhou, F., Caron, P., Mangeot, P.E., Ricci, E.P., Mourad, R., et al. (2021). Loop extrusion as a mechanism for formation of DNA damage repair foci. Nature.

Ba, Z., Lou, J., Ye, A.Y., Dai, H.Q., Dring, E.W., Lin, S.G., Jain, S., Kyritsis, N., Kieffer-Kwon, K.R., Casellas, R., et al. (2020). CTCF orchestrates long-range cohesin-driven V(D)J recombinational scanning. Nature 586, 305–310.

Banigan, E.J., van den Berg, A.A., Brandao, H.B., Marko, J.F., and Mirny, L.A. (2020). Chromosome organization by one-sided and two-sided loop extrusion. Elife 9.

Beckouet, F., Hu, B., Roig, M.B., Sutani, T., Komata, M., Uluocak, P., Katis, V.L., Shirahige, K., and Nasmyth, K. (2010). An Smc3 acetylation cycle is essential for establishment of sister chromatid cohesion. Molecular cell 39, 689–699.

Beckouet, F., Srinivasan, M., Roig, M.B., Chan, K.L., Scheinost, J.C., Batty, P., Hu, B., Petela, N., Gligoris, T., Smith, A.C., et al. (2016). Releasing Activity Disengages Cohesin’s Smc3/Scc1 Interface in a Process Blocked by Acetylation. Molecular cell 61, 563–574.

Chan, K.L., Gligoris, T., Upcher, W., Kato, Y., Shirahige, K., Nasmyth, K., and Beckouet, F. (2013). Pds5 promotes and protects cohesin acetylation. Proceedings of the National Academy of Sciences of the United States of America 110, 13020–13025.

Chan, K.L., Roig, M.B., Hu, B., Beckouet, F., Metson, J., and Nasmyth, K. (2012). Cohesin’s DNA Exit Gate Is Distinct from Its Entrance Gate and Is Regulated by Acetylation. Cell 150, 961–974.

Chapard, C., Jones, R., van Oepen, T., Scheinost, J.C., and Nasmyth, K. (2019). Sister DNA Entrapment between Juxtaposed Smc Heads and Kleisin of the Cohesin Complex. Molecular cell 75, 224–237 e225.

Cockram, C., Thierry, A., Gorlas, A., Lestini, R., and Koszul, R. (2021). Euryarchaeal genomes are folded into SMC-dependent loops and domains, but lack transcription-mediated compartmentalization. Molecular cell 81, 459–472 e410.

Collier, J.E., Lee, B.G., Roig, M.B., Yatskevich, S., Petela, N.J., Metson, J., Voulgaris, M., Gonzalez Llamazares, A., Lowe, J., and Nasmyth, K.A. (2020). Transport of DNA within cohesin involves clamping on top of engaged heads by Scc2 and entrapment within the ring by Scc3. Elife 9.

Costantino, L., Hsieh, T.S., Lamothe, R., Darzacq, X., and Koshland, D. (2020). Cohesin residency determines chromatin loop patterns. Elife 9.

Cournac, A., Marie-Nelly, H., Marbouty, M., Koszul, R., and Mozziconacci, J. (2012). Normalization of a chromosomal contact map. BMC Genomics 13, 436.

Dai, H.Q., Hu, H., Lou, J., Ye, A.Y., Ba, Z., Zhang, X., Zhang, Y., Zhao, L., Yoon, H.S., Chapdelaine-Williams, A.M., et al. (2021). Loop extrusion mediates physiological Igh locus contraction for RAG scanning. Nature.

Dauban, L., Montagne, R., Thierry, A., Lazar-Stefanita, L., Bastie, N., Gadal, O., Cournac, A., Koszul, R., and Beckouet, F. (2020). Regulation of Cohesin-Mediated Chromosome Folding by Eco1 and Other Partners. Molecular cell 77, 1279–1293 e1274.

Davidson, I.F., Bauer, B., Goetz, D., Tang, W., Wutz, G., and Peters, J.M. (2019). DNA loop extrusion by human cohesin. Science 366, 1338–1345.

Dekker, J., and Mirny, L. (2016). The 3D Genome as Moderator of Chromosomal Communication. Cell 164, 1110–1121.

Fernius, J., Nerusheva, O.O., Galander, S., Alves Fde, L., Rappsilber, J., and Marston, A.L. (2013). Cohesin-dependent association of scc2/4 with the centromere initiates pericentromeric cohesion establishment. Curr Biol 23, 599–606.

Ganji, M., Shaltiel, I.A., Bisht, S., Kim, E., Kalichava, A., Haering, C.H., and Dekker, C. (2018). Real-time imaging of DNA loop extrusion by condensin. Science 360, 102–105.

Gibcus, J.H., Samejima, K., Goloborodko, A., Samejima, I., Naumova, N., Nuebler, J., Kanemaki, M.T., Xie, L., Paulson, J.R., Earnshaw, W.C., et al. (2018). A pathway for mitotic chromosome formation. Science 359.

Goloborodko, A., Imakaev, M.V., Marko, J.F., and Mirny, L. (2016). Compaction and segregation of sister chromatids via active loop extrusion. Elife 5.

Gutierrez-Escribano, P., Newton, M.D., Llauro, A., Huber, J., Tanasie, L., Davy, J., Aly, I., Aramayo, R., Montoya, A., Kramer, H., et al. (2019). A conserved ATP- and Scc2/4-dependent activity for cohesin in tethering DNA molecules. Sci Adv 5, eaay6804.

Haarhuis, J.H.I., van der Weide, R.H., Blomen, V.A., Yanez-Cuna, J.O., Amendola, M., van Ruiten, M.S., Krijger, P.H.L., Teunissen, H., Medema, R.H., van Steensel, B., et al. (2017). The Cohesin Release Factor WAPL Restricts Chromatin Loop Extension. Cell 169, 693–707 e614.

Haering, C.H., Farcas, A.M., Arumugam, P., Metson, J., and Nasmyth, K. (2008). The cohesin ring concatenates sister DNA molecules. Nature 454, 297–301.

Higashi, T.L., Eickhoff, P., Sousa, J.S., Locke, J., Nans, A., Flynn, H.R., Snijders, A.P., Papageorgiou, G., O’Reilly, N., Chen, Z.A., et al. (2020). A Structure-Based Mechanism for DNA Entry into the Cohesin Ring. Molecular cell 79, 917–933 e919.

Hu, B., Petela, N., Kurze, A., Chan, K.L., Chapard, C., and Nasmyth, K. (2015). Biological chromodynamics: a general method for measuring protein occupancy across the genome by calibrating ChIP-seq. Nucleic Acids Res 43, e132.

Kikuchi, S., Borek, D.M., Otwinowski, Z., Tomchick, D.R., and Yu, H. (2016). Crystal structure of the cohesin loader Scc2 and insight into cohesinopathy. Proc Natl Acad Sci U S A 113, 12444–12449.

Kim, Y., Shi, Z., Zhang, H., Finkelstein, I.J., and Yu, H. (2019). Human cohesin compacts DNA by loop extrusion. Science 366, 1345–1349.

Kschonsak, M., Merkel, F., Bisht, S., Metz, J., Rybin, V., Hassler, M., and Haering, C.H. (2017). Structural Basis for a Safety-Belt Mechanism That Anchors Condensin to Chromosomes. Cell 171, 588–600 e524.

Lazar-Stefanita, L., Scolari, V.F., Mercy, G., Muller, H., Guerin, T.M., Thierry, A., Mozziconacci, J., and Koszul, R. (2017). Cohesins and condensins orchestrate the 4D dynamics of yeast chromosomes during the cell cycle. EMBO J 36, 2684–2697.

Le, T.B.K., Imakaev, M.V., Mirny, L.A., and Laub, M.T. (2013). High-Resolution Mapping of the Spatial Organization of a Bacterial Chromosome. Science 342, 731–734.

Lee, B.G., Roig, M.B., Jansma, M., Petela, N., Metson, J., Nasmyth, K., and Lowe, J. (2016). Crystal Structure of the Cohesin Gatekeeper Pds5 and in Complex with Kleisin Scc1. Cell Rep 14, 2108–2115.

Li, Y., Haarhuis, J.H.I., Sedeno Cacciatore, A., Oldenkamp, R., van Ruiten, M.S., Willems, L., Teunissen, H., Muir, K.W., de Wit, E., Rowland, B.D., et al. (2020). The structural basis for cohesin-CTCF-anchored loops. Nature 578, 472–476.

Makrantoni, V., and Marston, A.L. (2018). Cohesin and chromosome segregation. Curr Biol 28, R688–R693.

Marbouty, M., Le Gall, A., Cattoni, D.I., Cournac, A., Koh, A., Fiche, J.B., Mozziconacci, J., Murray, H., Koszul, R., and Nollmann, M. (2015). Condensin- and Replication-Mediated Bacterial Chromosome Folding and Origin Condensation Revealed by Hi-C and Superresolution Imaging. Molecular cell 59, 588–602.

Marchal, C., Sima, J., and Gilbert, D.M. (2019). Control of DNA replication timing in the 3D genome. Nat Rev Mol Cell Biol 20, 721–737.

Matthey-Doret, C., Baudry, L., Breuer, A., Montagne, R., Guiglielmoni, N., Scolari, V., Jean, E., Campeas, A., Chanut, P.H., Oriol, E., et al. (2020). Computer vision for pattern detection in chromosome contact maps. Nat Commun 11, 5795.

Merkenschlager, M., and Nora, E.P. (2016). CTCF and Cohesin in Genome Folding and Transcriptional Gene Regulation. Annu Rev Genom Hum G 17, 17–43.

Muir, K.W., Kschonsak, M., Li, Y., Metz, J., Haering, C.H., and Panne, D. (2016). Structure of the Pds5-Scc1 Complex and Implications for Cohesin Function. Cell Rep 14, 2116–2126.

Muller, H., Scolari, V.F., Agier, N., Piazza, A., Thierry, A., Mercy, G., Descorps-Declere, S., Lazar-Stefanita, L., Espeli, O., Llorente, B., et al. (2018). Characterizing meiotic chromosomes’ structure and pairing using a designer sequence optimized for Hi-C. Mol Syst Biol 14, e8293.

Nasmyth, K. (2001). Disseminating the genome: joining, resolving, and separating sister chromatids during mitosis and meiosis. Annual review of genetics 35, 673–745.

Nasmyth, K., and Haering, C.H. (2009). Cohesin: Its Roles and Mechanisms. Annual review of genetics 43, 525–528.

Nora, E.P., Lajoie, B.R., Schulz, E.G., Giorgetti, L., Okamoto, I., Servant, N., Piolot, T., van Berkum, N.L., Meisig, J., Sedat, J., et al. (2012). Spatial partitioning of the regulatory landscape of the X-inactivation centre. Nature 485, 381–385.

Ouyang, Z., Zheng, G., Tomchick, D.R., Luo, X., and Yu, H. (2016). Structural Basis and IP6 Requirement for Pds5-Dependent Cohesin Dynamics. Molecular cell 62, 248–259.

Paldi, F., Alver, B., Robertson, D., Schalbetter, S.A., Kerr, A., Kelly, D.A., Baxter, J., Neale, M.J., and Marston, A.L. (2020). Convergent genes shape budding yeast pericentromeres. Nature 582, 119–123.

Petela, N.J., Gligoris, T.G., Metson, J., Lee, B.G., Voulgaris, M., Hu, B., Kikuchi, S., Chapard, C., Chen, W., Rajendra, E., et al. (2018). Scc2 Is a Potent Activator of Cohesin’s ATPase that Promotes Loading by Binding Scc1 without Pds5. Molecular cell 70, 1134–1148 e1137.

Piazza, A., Bordelet, H., Dumont, A., Thierry, A., Savocco, J., Girard, F., and Koszul, R. (2020). Cohesin regulates homology search during recombinational DNA repair. bioRxiv, 2020.2012.2017.423195.

Rao, S.S., Huntley, M.H., Durand, N.C., Stamenova, E.K., Bochkov, I.D., Robinson, J.T., Sanborn, A.L., Machol, I., Omer, A.D., Lander, E.S., et al. (2014). A 3D map of the human genome at kilobase resolution reveals principles of chromatin looping. Cell 159, 1665–1680.

Rao, S.S.P., Huang, S.C., Glenn St Hilaire, B., Engreitz, J.M., Perez, E.M., Kieffer-Kwon, K.R., Sanborn, A.L., Johnstone, S.E., Bascom, G.D., Bochkov, I.D., et al. (2017). Cohesin Loss Eliminates All Loop Domains. Cell 171, 305–320 e324.

Rolef Ben-Shahar, T., Heeger, S., Lehane, C., East, P., Flynn, H., Skehel, M., and Uhlmann, F. (2008). Eco1-dependent cohesin acetylation during establishment of sister chromatid cohesion. Science 321, 563–566.

Rowland, B.D., Roig, M.B., Nishino, T., Kurze, A., Uluocak, P., Mishra, A., Beckouet, F., Underwood, P., Metson, J., Imre, R., et al. (2009). Building sister chromatid cohesion: smc3 acetylation counteracts an antiestablishment activity. Molecular cell 33, 763–774.

Rowley, M.J., and Corces, V.G. (2018). Organizational principles of 3D genome architecture. Nat Rev Genet 19, 789–800.

Sanborn, A.L., Rao, S.S., Huang, S.C., Durand, N.C., Huntley, M.H., Jewett, A.I., Bochkov, I.D., Chinnappan, D., Cutkosky, A., Li, J., et al. (2015). Chromatin extrusion explains key features of loop and domain formation in wild-type and engineered genomes. Proc Natl Acad Sci U S A 112, E6456–6465.

Schalbetter, S.A., Goloborodko, A., Fudenberg, G., Belton, J.M., Miles, C., Yu, M., Dekker, J., Mirny, L., and Baxter, J. (2017). SMC complexes differentially compact mitotic chromosomes according to genomic context. Nat Cell Biol 19, 1071–1080.

Schwarzer, W., Abdennur, N., Goloborodko, A., Pekowska, A., Fudenberg, G., Loe-Mie, Y., Fonseca, N.A., Huber, W., C, H.H., Mirny, L., et al. (2017). Two independent modes of chromatin organization revealed by cohesin removal. Nature 551, 51–56.

Shi, Z., Gao, H., Bai, X.C., and Yu, H. (2020). Cryo-EM structure of the human cohesin-NIPBL-DNA complex. Science 368, 1454–1459.

Srinivasan, M., Petela, N.J., Scheinost, J.C., Collier, J., Voulgaris, M., M, B.R., Beckouet, F., Hu, B., and Nasmyth, K.A. (2019). Scc2 counteracts a Wapl-independent mechanism that releases cohesin from chromosomes during G1. Elife 8.

Szabo, Q., Bantignies, F., and Cavalli, G. (2019). Principles of genome folding into topologically associating domains. Science Advances 5, eaaw1668.

Unal, E., Heidinger-Pauli, J.M., Kim, W., Guacci, V., Onn, I., Gygi, S.P., and Koshland, D.E. (2008). A molecular determinant for the establishment of sister chromatid cohesion. Science 321, 566–569.

Wutz, G., Ladurner, R., St Hilaire, B.G., Stocsits, R.R., Nagasaka, K., Pignard, B., Sanborn, A., Tang, W., Varnai, C., Ivanov, M.P., et al. (2020). ESCO1 and CTCF enable formation of long chromatin loops by protecting cohesin(STAG1) from WAPL. Elife 9.

Wutz, G., Varnai, C., Nagasaka, K., Cisneros, D.A., Stocsits, R.R., Tang, W., Schoenfelder, S., Jessberger, G., Muhar, M., Hossain, M.J., et al. (2017). Topologically associating domains and chromatin loops depend on cohesin and are regulated by CTCF, WAPL, and PDS5 proteins. EMBO J 36, 3573–3599.

Yatskevich, S., Rhodes, J., and Nasmyth, K. (2019). Organization of Chromosomal DNA by SMC Complexes. Annual review of genetics 53, 445–482.

Yu, M., and Ren, B. (2017). The Three-Dimensional Organization of Mammalian Genomes. Annu Rev Cell Dev Biol 33, 265–289.

Zhang, J., Shi, X., Li, Y., Kim, B.J., Jia, J., Huang, Z., Yang, T., Fu, X., Jung, S.Y., Wang, Y., et al. (2008). Acetylation of Smc3 by Eco1 is required for S phase sister chromatid cohesion in both human and yeast. Molecular cell 31, 143–151.

Zhang, X., Zhang, Y., Ba, Z., Kyritsis, N., Casellas, R., and Alt, F.W. (2019a). Fundamental roles of chromatin loop extrusion in antibody class switching. Nature 575, 385–389.

Zhang, Y., Zhang, X., Ba, Z., Liang, Z., Dring, E.W., Hu, H., Lou, J., Kyritsis, N., Zurita, J., Shamim, M.S., et al. (2019b). The fundamental role of chromatin loop extrusion in physiological V(D)J recombination. Nature 573, 600–604.

Zheng, H., and Xie, W. (2019). The role of 3D genome organization in development and cell differentiation. Nat Rev Mol Cell Biol 20, 535–550.

